# Understanding the source of METTL3-independent m^6^A in mRNA

**DOI:** 10.1101/2021.12.15.472866

**Authors:** Hui Xian Poh, Aashiq H. Mirza, Brian F. Pickering, Samie R. Jaffrey

## Abstract

*N*^6^-methyladenosine (m^6^A) is a highly prevalent mRNA modification which promotes degradation of transcripts encoding proteins that have roles in cell development, differentiation, and other pathways. METTL3 is the major methyltransferase that catalyzes the formation of m^6^A in mRNA. As 30–80% of m^6^A can remain in mRNA after METTL3 depletion by CRISPR/Cas9-based methods, other enzymes are thought to catalyze a sizable fraction of m^6^A. Here, we re-examined the source of m^6^A in the mRNA transcriptome. We characterized mouse embryonic stem cell lines which continue to have m^6^A in their mRNA after *Mettl3* knockout. We show that these cells express alternatively spliced *Mettl3* transcript isoforms that bypass the CRISPR/Cas9 mutations and produce functionally active methyltransferases. We similarly show that other reported *METTL3* knockout cell lines express altered METTL3 proteins. We find that gene dependency datasets show that most cell lines fail to proliferate after *METTL3* deletion, suggesting that reported *METTL3* knockout cell lines express altered METTL3 proteins rather than have full knockout. Finally, we reassessed METTL3’s role in synthesizing m^6^A using a genomic deletion of *Mettl3*, and found that METTL3 is responsible for >95% of m^6^A in mRNA. Overall, these studies suggest that METTL3 is responsible for the vast majority of m^6^A in the transcriptome, and that remaining m^6^A in putative *METTL3* knockout cell lines is due to the expression of altered but functional METTL3 isoforms.

## INTRODUCTION

*N*^6^-methyladenosine (m^6^A) is the most abundant internal mRNA modification [1–3] and its presence in mRNA is critical for cellular differentiation [4–6], cancer progression [7–9], and other cellular processes [10–14]. m^6^A in mRNA can regulate mRNA fates (reviewed by [15–17]), particularly by promoting mRNA degradation [18,19].

The first enzyme shown to catalyze m^6^A formation was METTL3 [18], which forms a heterodimer complex with METTL14 [19–21]. METTL3 contains the catalytic component. METTL14 was initially believed to have catalytic ability [21], but METTL14 is now known to be catalytically inactive [19,20]. Instead, METTL14 binds and positions RNA for methylation [19,20]. The METTL3-METTL14 complex is a component of a larger multiprotein “m^6^A writer complex” that mediates co-transcriptional mRNA methylation [1,2,22].

Although METTL3 is often described as the major m^6^A-forming enzyme in cells, the amount of m^6^A thought to be formed by METTL3 varies widely in different studies. The first study to knockout *Mettl3* showed that ~40% of m^6^A remained after *Mettl3* knockout in mouse embryonic stem cells (mESCs) [4]. The authors suggested that METTL14 may account for this 40% of m^6^A, based on the previous understanding that METTL14 was catalytic. However, a different group shortly thereafter reported that deletion of either *Mettl3* or *Mettl14* in mESCs leads to a loss of ~99% of m^6^A in mRNA [5]. These contradictory results have led to uncertainty about how much m^6^A in mRNA derives from METTL3.

*METTL3* has been knocked out in other cell lines and tissues. These results have shown that 30–80% of m^6^A can remain after *METTL3* knockout [23–33]. In U2OS osteosarcoma cells, ~60% of m^6^A remained after CRISPR-mediated knockout of *METTL3* [23,24]. In HEK293T human embryonic kidney cells, ~50% of m^6^A remained after CRISPR-mediated knockout of *METTL3* [25]. After Cre-conditional genomic deletion of *Mettl3* in mouse CD4+ T-cells, 28% of m^6^A remained [30]. Since a non-negligible amount of m^6^A persists after *METTL3* knockout, it has been speculated that other methyltransferases may have a major role in forming m^6^A in mRNA [4,31,33,34].

Here, we address the source of m^6^A in mRNA in reported *METTL3* knockout cells. We examined two different *Mettl3* knockout mESC lines, which both report loss of METTL3, but showed vastly different levels of residual m^6^A in mRNA. We show that *Mettl3* mutagenesis by CRISPR approaches can create alternatively spliced isoforms of *Mettl3*, resulting in an altered but catalytically active METTL3 protein. Thus, the residual m^6^A can be ascribed to a hypomorphic *METTL3* allele. We further show that other published *METTL3* mutant cell lines, which were intended to delete *METTL3*, retain m^6^A and express alternative METTL3 proteins. Furthermore, we show that *METTL3* is an essential gene in most cell lines, and thus, *METTL3* knockout cell lines which remain viable are likely to have generated alternatively spliced functional METTL3 proteins that bypass the CRISPR mutations. Lastly, we show that when genomic deletion is used to create a large deletion of *METTL3*, essentially all m^6^A is depleted in a fibroblast cell line. Overall, these studies argue that METTL3 is responsible for most m^6^A in mRNA, and that residual m^6^A after *METTL3* depletion usually reflects the generation of hypomorphic *METTL3* alleles and therefore incomplete *METTL3* knockout.

## RESULTS

### Two mESC lines exhibit different levels of m^6^A after *Mettl3* knockout

To understand how much m^6^A in mRNA is catalyzed by METTL3, we examined two previously reported *Mettl3* knockout mESC lines. Two groups independently knocked out *Mettl3* in mESCs and reported markedly different levels of residual m^6^A levels in mRNA [4,5]. The first *Mettl3* knockout mESC line was described in Batista et al. by Howard Chang’s group, and used a CRISPR/Cas9 approach [4]. The guide RNAs were designed to introduce deletions in exon 2 of *Mettl3* and create premature termination codons [4]. The resulting mESC line, designated “HC *Mettl3* KO mESCs”, was found to have 40% residual m^6^A in mRNA. These authors understandably attributed the remaining m^6^A to METTL14 since, at that time, METTL14 was incorrectly shown to be a functional methyltransferase [21]. The second mESC line was described in Geula *et al* by Jacob Hanna’s group [5]. This group used *loxP* sites surrounding exon 4 in *Mettl3* to delete the exon encoding the zinc finger domain (ZFD), an RNA recognition domain required for METTL3 methyltransferase activity [19,35]. This *Mettl3* knockout mESC line, designated “JH *Mettl3* KO mESCs,” exhibited <1% remaining m^6^A in mRNA. It is unclear why the HC *Mettl3* KO mESCs have high m^6^A levels when the JH *Mettl3* KO mESCs, which in principle should be the same, have virtually no remaining m^6^A.

We first re-confirmed the levels of m^6^A in these two cell lines. The m^6^A levels in the two *Mettl3* knockout mESC lines were originally measured via different methods, which may have led to these contradictory results. Thin-layer chromatography was used by Batista *et al*, while mass spectrometry was used by Geula *et al*. We measured m^6^A in the mESC lines using mass spectrometry [36]. Our mass spectrometry measurements were consistent with the results originally reported by the two groups. In the two HC *Mettl3* KO mESC lines, designated *Mettl3*-KO-mESC-a and *Mettl3*-KO-mESC-b, we saw 40.2% and 55.6% residual m^6^A (**Fig 1A**), comparable to ~40% originally reported by this group [4]. However, the JH *Mettl3* KO mESCs had only 1.45% residual m^6^A (**Fig 1A**), corroborating the 0.5% remaining m^6^A originally reported by this group [5].

**Fig 1.**
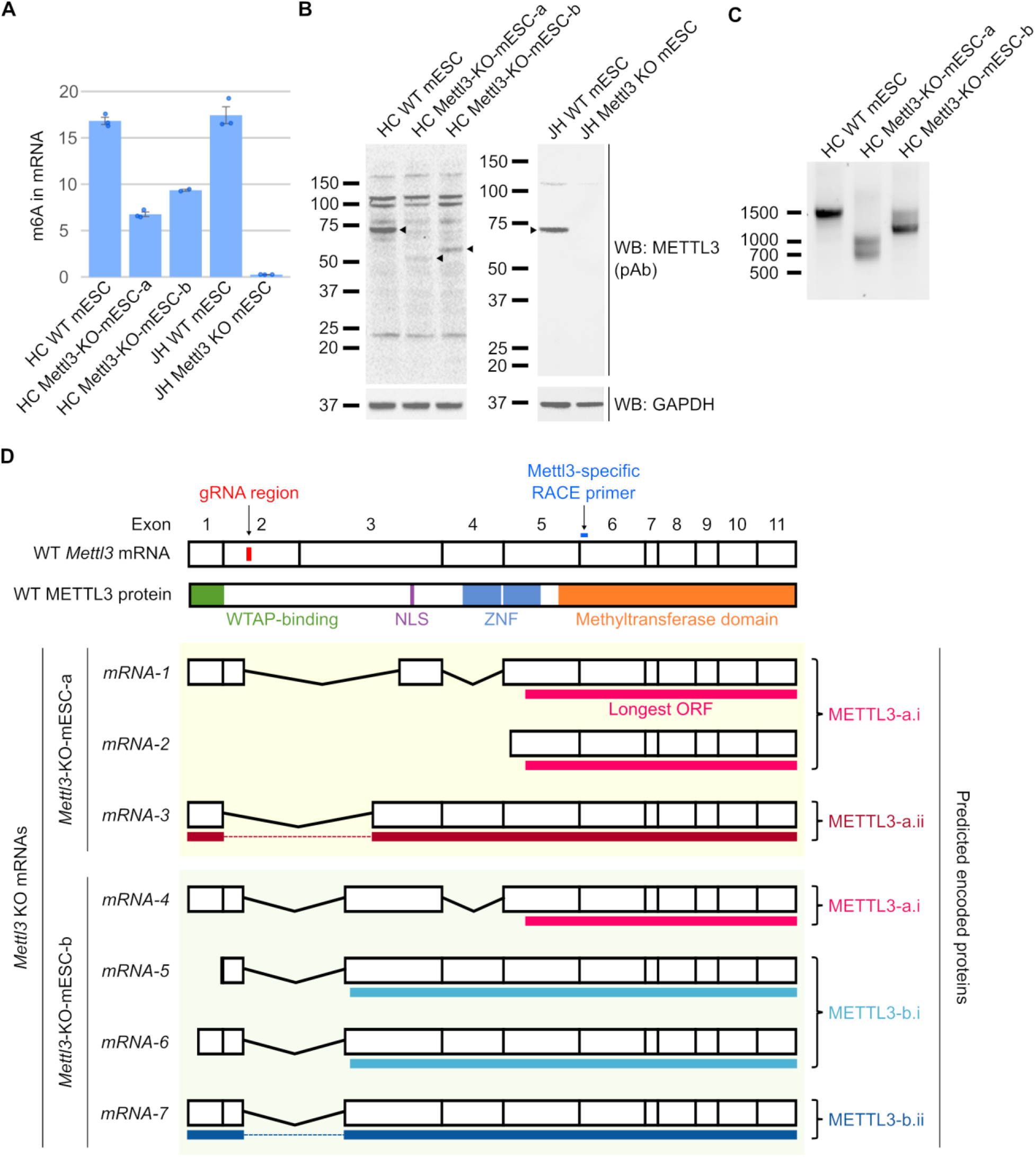
Previously described *Mettl3* KO mESC lines express shorter isoforms of *Mettl3*. **A** *Mettl3* KO mESCs from two groups have different m^6^A levels. To re-confirm the m^6^A levels using quantitative methods, we performed mass spectrometry to estimate the m^6^A in mRNA. HC *Mettl3* KO mESCs show persistence of 40.2% (*Mettl3*-KO-mESC-a) and 55.6% (*Mettl3*-KO-mESC-b) m^6^A respectively, while JH *Mettl3* KO mESCs show 1.45% m^6^A compared to wild-type. This confirms that JH *Mettl3* KO mESCs have near-complete loss of m^6^A, but not the HC *Mettl3* KO mESCs. Error bars indicate standard error (n = 3 for all, except n = 2 for HC *Mettl3*-KO-mESC-b). **B** HC *Mettl3* KO mESCs exhibit new anti-METTL3-immunoreactive bands. To investigate the effectiveness of the *Mettl3* KO, we measured the loss of METTL3 via western blot. Full-length METTL3 (75 kDa, arrowhead) was lost in both KO cell lines, but new bands, which were reactive to the anti-METTL3-antibody, appeared at ~50 kDa in *Mettl3*-KO-mESC-a and ~55 kDa in *Mettl3*-KO-mESC-b (arrowheads). This indicates the possibility that a novel smaller METTL3 protein was expressed in the KO mESCs. In contrast, JH *Mettl3* KO mESCs have no proteins reactive to anti-METTL3-antibodies. 30 µg per lane. WB = western blot, pAb = polyclonal antibody. **C** 5’ RACE reveals the expression of shorter *Mettl3* mRNAs in the *Mettl3* KO mESCs. We used 5’ RACE to identify novel *Mettl3* mRNAs in the *Mettl3* KO mESCs. The full-length RACE product (~1500 bp) was lost in the KO cells, but novel products at ~1000 bp and ~700 bp were found in *Mettl3*-KO-mESC-a and at ~1500 bp and ~1300 bp in *Mettl3*-KO-mESC-b. These shorter mRNAs may encode the smaller METTL3 proteins seen in the KO cells. **D** Sequencing of 5’ RACE products show *Mettl3* mRNAs with exon skipping or alternative transcription-start sites. We sequenced the 5’ RACE products to characterize the *Mettl3* mRNA transcripts that are expressed by the *Mettl3* KO mESCs. All *Mettl3* mRNAs expressed in the KO cells skipped the guide RNA deletion region by exon skipping, or by using alternative transcription-start sites downstream of the deletion. The longest ORFs which are in-frame with the wild-type *Mettl3* mRNAs are shown as solid lines below each mRNA. The encoded protein is also represented, with the domains required for METTL3 activity shown. ZFD = zinc finger domain; NLS = nuclear localization signal.

The near-complete loss of m^6^A in the JH *Mettl3* KO mESCs suggests that METTL3 is the major m^6^A writer in this mESC line. On the other hand, the HC *Mettl3* KO mESCs still retain m^6^A despite *Mettl3* depletion. Although it is possible that these mESCs use an alternate enzyme for m^6^A biosynthesis, we suspected that *Mettl3* was not completely knocked out in these cell lines.

### *Mettl3* knockout mESCs which retain m^6^A express alternative METTL3 isoforms

We next wanted to confirm that *Mettl3* was knocked out in the HC *Mettl3* KO mESCs. Previously, a western blot was used to determine the loss of METTL3 protein [4]. To first confirm that the METTL3 protein is indeed absent, we performed a western blot using a METTL3 polyclonal antibody. Full-length METTL3 (~75 kDa) was identified in the wild-type (WT) mESCs, and was lost in both *Mettl3* KO cell lines (**Fig 1B**). However, we observed new bands in the anti-METTL3 immunoblot in the knockout cell lines. The new proteins were ~50 kDa in *Mettl3*-KO-mESC-a, and ~55 kDa in *Mettl3*-KO-mESC-b (**Fig 1B**). These proteins were not visible in the wild-type cell line, suggesting they are unique to the knockout cell lines. These proteins were also not visible in the JH *Mettl3* KO cell lines (**Fig 1B**). To confirm that these are indeed METTL3 proteins, we repeated the western blot with a second anti-METTL3 antibody. Again, we found the same bands in the *Mettl3* KO cell lines (**Fig S1A, S1B**). These proteins may have escaped notice in the original study due to the smaller mass and lower expression of these bands compared to wild-type METTL3. Overall, the new METTL3-antibody-reactive proteins suggests that smaller METTL3 isoforms are produced in the *Mettl3* KO cells which may be the source of m^6^A in these cells.

We next wanted to determine if these smaller METTL3 proteins are functional and could account for the m^6^A produced in the HC *Mettl3* KO cells. To do this, we first determined the sequence of these isoforms. Since the HC *Mettl3* KO cells were produced using guide RNAs targeting exon 2 of *Mettl3* [4], any CRISPR/Cas9-induced mutations and potential alternative splicing events are likely to be in the 5’ end of the transcript. We therefore used 5’ RACE (rapid amplification of cDNA ends) [37,38] to identify new transcription-start sites or possible exon skipping events of the *Mettl3* transcripts expressed in these cells.

Using 5’ RACE, a single major ~1500 bp band was seen for *Mettl3* in wild-type mESCs (**Fig 1C**). In contrast, in the two HC *Mettl3* KO cell lines, we found shorter bands indicative of *Mettl3* mRNAs with a shorter 5’ region (**Fig 1C**). We sequenced these 5’ RACE products and found several *Mettl3* mRNAs from the *Mettl3* KO cells (**Fig 1D, S1 Table**). In all cases, the mRNAs show alternative splicing that bypasses the CRISPR deletion in exon 2 by exon skipping, or use of an alternative transcription-start site downstream of the deletion.

We next asked if these mRNAs could potentially encode the altered METTL3 proteins detected in the *Mettl3* KO mESCs. For each mRNA, we identified the longest possible ORF which is in frame with the METTL3 catalytic domain (**Fig 1D**). In the case of *Mettl3*-KO-mESC-a, we found a transcript that is predicted to encode a METTL3 protein (designated “METTL3-a.ii”) that matches the size of the altered METTL3 protein from this cell line (**Fig 1B, 1D**). Similarly, we identified a transcript that is predicted to encode the altered METTL3 protein in *Mettl3*-KO-mESC-b (designated “METTL3-b.ii”) (**Fig 1B, 1D**).

Together, these data identify potential transcripts that may encode the altered METTL3 proteins found in the HC *Mettl3* KO cells.

### Altered METTL3 proteins expressed in incomplete *Mettl3* knockout mESCs can catalyze the formation of m^6^A in cells

We next asked if the novel *Mettl3* transcript isoforms encode functional m^6^A-forming methyltransferases. METTL3 contains several domains necessary for methyltransferase activity, including the WTAP-binding domain [37], the zinc finger domain (ZFD) [19,35], and the methyltransferase domain [19,20]. Both METTL3-a.ii and METTL3-b.ii contain all three domains, and therefore may be functional. Other altered *Mettl3* transcripts in the *Mettl3* KO mESCs encode predicted proteins (designated “METTL3-a.i” and “METTL3-b.i”) that lack one or more of these domains (**Fig 2A**).

**Fig 2.**
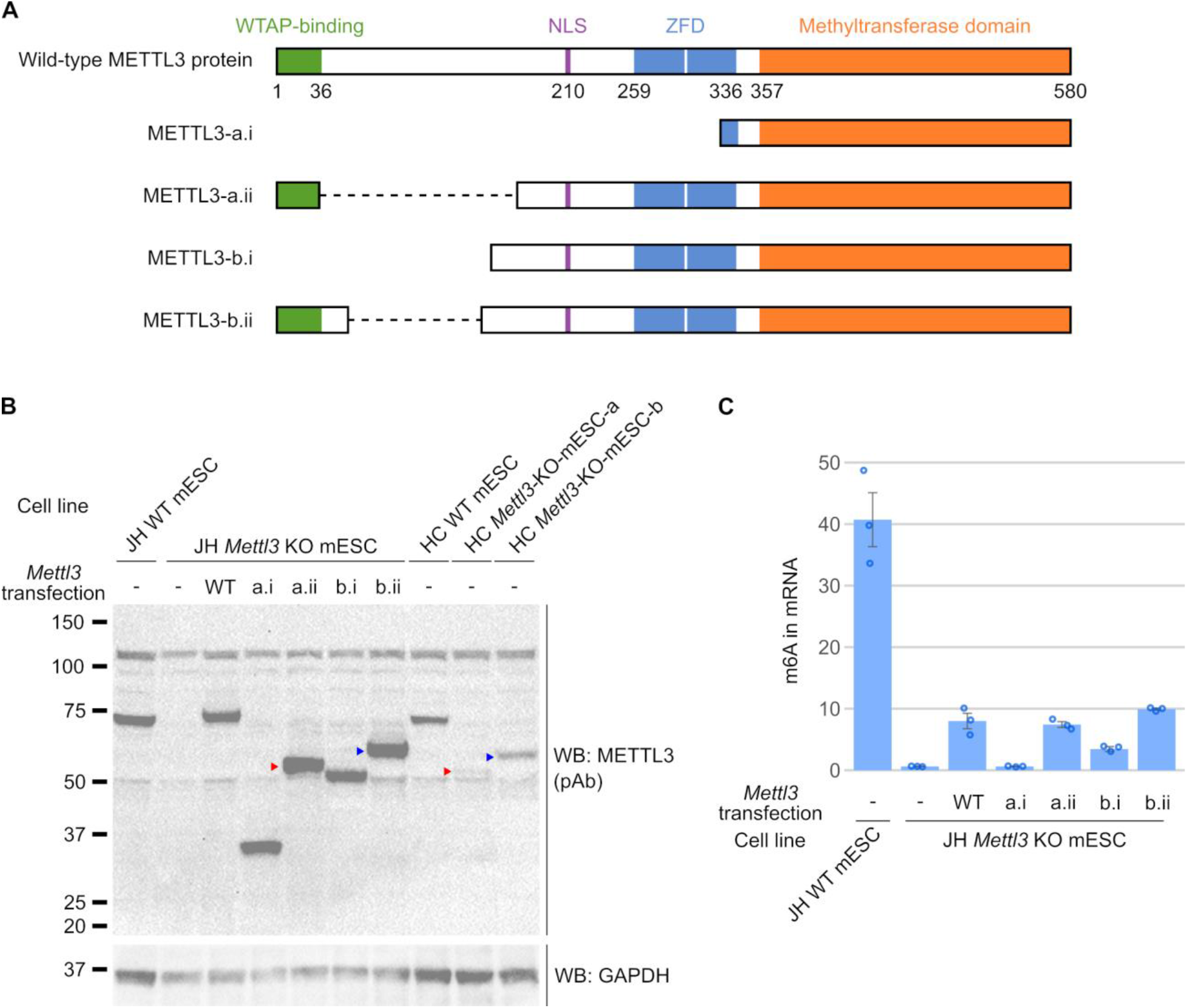
Shortened isoforms of METTL3 catalyze m^6^A formation in *Mettl3* KO mESCs. **A** The predicted domain structure of proteins encoded by the altered *Mettl3* ORFs expressed in HC *Mettl3* KO mESCs suggest they may be functional. The domains that are known to be necessary for m^6^A formation by METTL3 include the WTAP-binding domain [37], ZFD (zinc finger domain) [19,35], and methyltransferase domains [19,20]. To determine if the *Mettl3* ORFs from *Mettl3* KO mESCs encode functional METTL3 proteins, we predicted the domain structure of the METTL3 protein isoforms from their ORFs. While all the predicted proteins have the SAM-binding and METTL14-binding domains, only METTL3-a.ii and METTL3-b.ii have all the known critical domains for m^6^A-writing. ZFD = zinc finger domain; NLS = nuclear localization signal. **B** Western blot of transfected *Mettl3* isoform ORFs. To determine if the *Mettl3* ORFs we found in the KO cells can synthesize m^6^A, we expressed the *Mettl3* ORFs in JH *Mettl3* KO mESCs which exhibit no METTL3 protein and essentially no baseline m^6^A signal. After 48 hours, the alternatively spliced METTL3 proteins can be detected by immunoblotting with an anti-METTL3 antibody. METTL3-a.ii (50kDa, red arrowhead) and METTL3-b.ii (55kDa, blue arrowhead) have similar sizes to the anti-METTL3-antibody-reactive protein seen in HC *Mettl3*-KO-mESC-a (red arrowhead) and *Mettl3*-KO-mESC-b (blue arrowhead), respectively. 30 µg per lane. WB = western blot, pAb = polyclonal antibody. **C** Isoforms of METTL3 proteins can write m^6^A. After 48 hours of transfection, mRNA from each sample was processed and m^6^A was measured using mass spectrometry. Expression of full-length wild-type METTL3 was able to rescue 19.7% of the m^6^A. METTL3-a.i was unable to rescue m^6^A, but METTL3-a.ii was able to rescue 18.3% of m^6^A. METTL3-b.i could only rescue 8.5% of m^6^A, whereas METTL3-b.ii rescued 24.4% of the m^6^A respectively. Thus, METTL3-a.ii and METTL3-b.ii proteins which are expressed in the HC *Mettl3* KO mESCs are able to catalyze the formation of m^6^A. Error bars indicate standard error (n = 3).

To test the activity of these METTL3 isoforms, we expressed them in the JH *Mettl3* KO mESCs, which lack METTL3-immunoreactive bands (**Fig 1B, Fig S1B**) and lack m^6^A in mRNA (**Fig 1A**). First, we noted that the size of METTL3-a.ii and METTL3-b.ii were indeed similar to the sizes of the smaller METTL3 proteins in *Mettl3*-KO-mESC-a and *Mettl3*-KO-mESC-b respectively (**Fig 2B**). Thus, METTL3-a.ii and METTL3-b.ii may be the novel METTL3 isoforms we detected in the HC *Mettl3* KO mESCs. In contrast, bands corresponding to METTL3-a.i (~30.4 kDa) and METTL3-b.i (~49 kDa) were not seen in *Mettl3*-KO-mESC-a or *Mettl3*-KO-mESC-b, respectively (**Fig 1B, 2B**). These METTL3 isoforms may have been less efficiently transcribed or translated, or may be less stable, leading to the lack of an immunoreactive band for these proteins.

We first tested expression of full-length METTL3 in the JH *Mettl3* KO mESCs, and found that they rescued m^6^A levels to 19.7% of wild-type (**Fig 2C**). The lack of complete rescue by the wild-type *Mettl3* transcript likely reflects the short time of expression (48 hours). METTL3-a.ii and METTL3-b.ii were able to rescue 18.3% and 24.4% of the m^6^A respectively (**Fig 2C**), suggesting that they are functional m^6^A methyltransferases. As predicted, METTL3-a.i failed to rescue m^6^A and METTL3-b.i could only rescue a small portion of m^6^A (8.5%) (**Fig 2C**). Overall, these data indicate that the HC *Mettl3* knockout mESCs express hypomorphic *Mettl3* alleles which encode catalytically active METTL3 isoforms i.e. METTL3-a.ii and METTL3-b.ii. Thus, the HC *Mettl3* knockout mESCs should be viewed as incomplete knockouts with hypomorphic *Mettl3* alleles.

### *METTL3* knockout U2OS cells express an altered METTL3 protein

The idea that METTL3 is not responsible for all m^6^A in mRNA was further supported by *METTL3* knockout in several cell lines, each of which show high residual levels of m^6^A [23–29]. For example, a *METTL3* KO U2OS cell line was reported to have ~60% of the m^6^A remaining compared to wild-type [23,24]. Similarly, we found that a previously reported *METTL3* KO A549 cell line [38] has ~90% of m^6^A remaining compared to the wild-type (**Fig S2A**). The persistence of m^6^A in these *METTL3* knockout cell lines has led to the idea that other enzymes mediate a substantial fraction of m^6^A in mRNA. However, in light of our finding that alternatively spliced *Mettl3* isoforms were induced in the *Mettl3* knockout mESCs, another possibility is that *METTL3* isoforms were induced in these *METTL3* knockout cell lines as well, which led to the high m^6^A levels in these *METTL3* knockout cell lines.

We examined one of these cell lines—the *METTL3* knockout U2OS cell line which was generated using CRISPR/Cas9-directed mutagenesis with a guide RNA targeting exon 1 [23]. We first reconfirmed the m^6^A levels in these *METTL3* knockout cells, and found that the *METTL3* knockout U2OS cells retained 75% m^6^A compared to the wild-type (**Fig 3A**), comparable to the 60% originally reported [24]. Hence, *METTL3* knockout U2OS cells continue to have high levels of m^6^A in mRNA.

**Fig 3.**
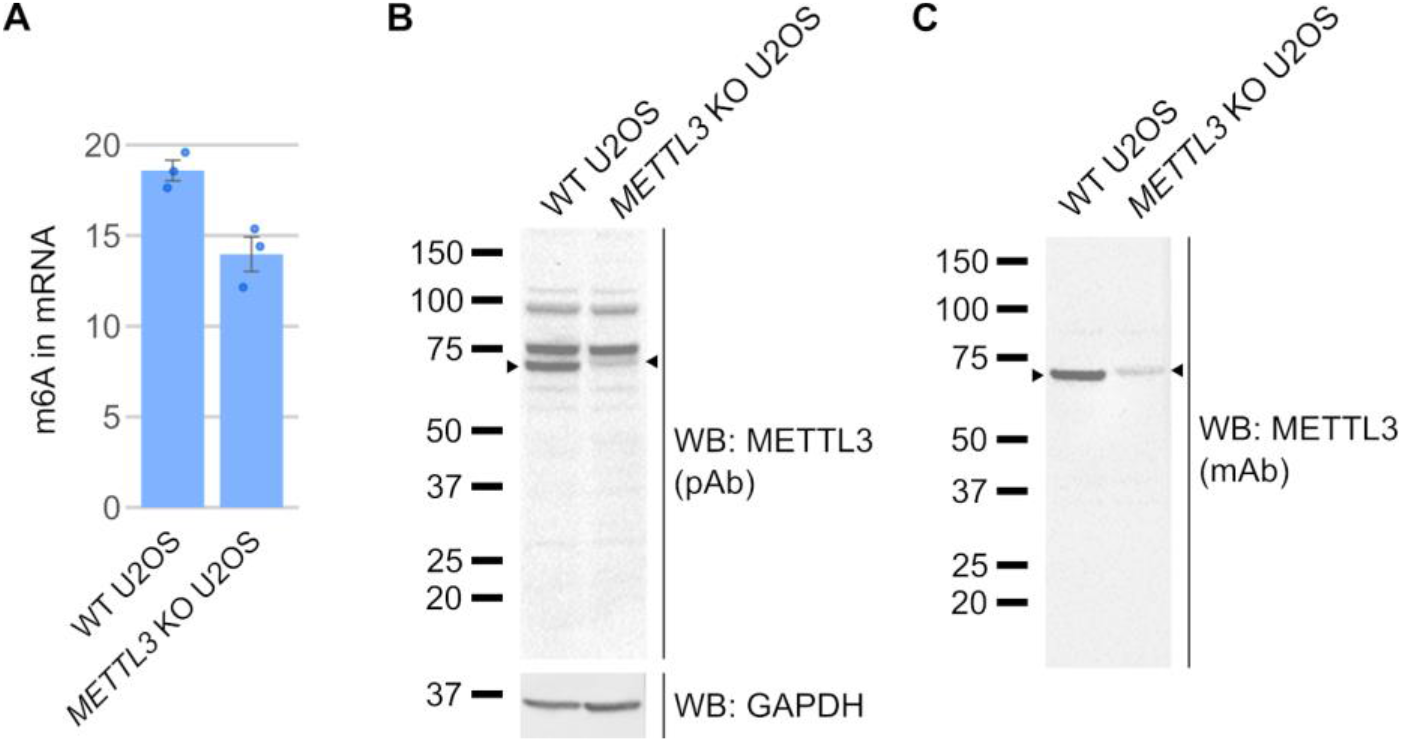
*METTL3* knockout in U2OS cells also appears to be incomplete. **A** *METTL3* KO U2OS cells have persistent m^6^A. *METTL3* KO U2OS cells have been reported to have 60% the levels of m^6^A found in control U2OS cells [24]. We re-confirmed this with mass spectrometry measurements of m^6^A, which showed that *METTL3* KO U2OS cells have 75.2% remaining m^6^A compared to wild-type. Thus, m^6^A levels remain high in *METTL3* KO U2OS cells. Error bars indicate standard error (n = 3). **B** *METTL3* KO U2OS cells express a novel anti-METTL3-antibody-reactive protein. As m^6^A levels were not completely ablated in the *METTL3* KO U2OS cells, we assessed METTL3 protein expression in these cells to confirm the if the knockout was effective. We found that wild-type METTL3 protein was lost in the *METTL3* KO U2OS cells, but a larger protein which was reactive to the anti-METTL3 antibody was found in the *METTL3* KO U2OS cells. This suggests the *METTL3* KO U2OS cells express a novel METTL3 protein that is slightly larger than the wild-type METTL3. WB = western blot, pAb = polyclonal antibody. **C** Confirmation of a novel METTL3-like protein in *METTL3* KO U2OS using a second METTL3 antibody. To confirm that the METTL3-immunoreactive band we saw in *METTL3* KO U2OS cells in **Fig 3B** was METTL3, we used a second anti-METTL3 monoclonal antibody to confirm the result. The same protein band is immunoreactive to the second anti-METTL3 antibody, thus suggesting that the *METTL3* KO U2OS cells express a novel, larger METTL3 protein. WB = western blot, mAb = monoclonal antibody.

We next asked if the *METTL3* KO U2OS cells express a METTL3 protein using a western blot. We observed full-length METTL3 in the wild-type U2OS cells, but not in the knockout cells (**Fig 3B**). However, a slightly larger anti-METTL3 immunoreactive band was detected exclusively in the knockout cells (**Fig 3B**). To confirm that this is indeed a METTL3 isoform, we validated the western blot with a second anti-METTL3 monoclonal antibody and found that the protein in the knockout cells was also reactive to the second anti-METTL3 antibody (**Fig 3C**). We note that we could observe a similar protein band in the authors’ original western blot which used a different METTL3 antibody [23], however it was much fainter and could easily be mistaken for non-specific background. Overall, this data again suggests that CRISPR/Cas9 mutagenesis results in the appearance of a novel METTL3 isoform in a *METTL3* knockout cell line, which could explain m^6^A persistence in these cells.

### *METTL3* knockout cell lines are generally not viable

Several *METTL3* knockout cell lines have been reported despite the fact that *METTL3* is thought to be an essential gene. *Mettl3* knockout is embryonic lethal in mice at E5.5, before cell specification occurs [5] and CRISPR screens have indicated that *METTL3* is an essential gene in specific cell lines that were tested [7,39]. On the other hand, the JH *Mettl3* knockout mESCs are able to survive, demonstrating that some cell lines can survive without *METTL3* or m^6^A. Therefore, it is not clear which cell lines require METTL3 for survival. If *METTL3* is required for survival of most cell lines, it would cast doubt on the stable *METTL3* knockout cell lines which have been reported in the literature.

To understand which cell lines require *METTL3* for survival, we screened the Cancer Dependency Map Project (DepMap) 21Q4 dataset [40–44]. This dataset measures cell proliferation in 1054 cell lines following a CRISPR loss-of-function screen. Failure of a cell line to grow after expression of the guide RNA which inactivates a gene indicates that the gene is essential in the cell line. The probability that a cell line is dependent on each gene is calculated as a gene dependency probability score, which accounts for guide efficacy as well as copy number of each gene. The 21Q4 DepMap dataset showed that *METTL3* is necessary for cell proliferation in 801 of 1054 tested cell lines (**Fig 4A**). The *METTL3*-dependent cell lines include the U2OS and A549 cell lines which other groups have used to make *METTL3* knockout cell lines [23,38] (**Fig 4A**). The DepMap data suggests that these reported *METTL3* cell lines should not have been viable.

**Fig 4.**
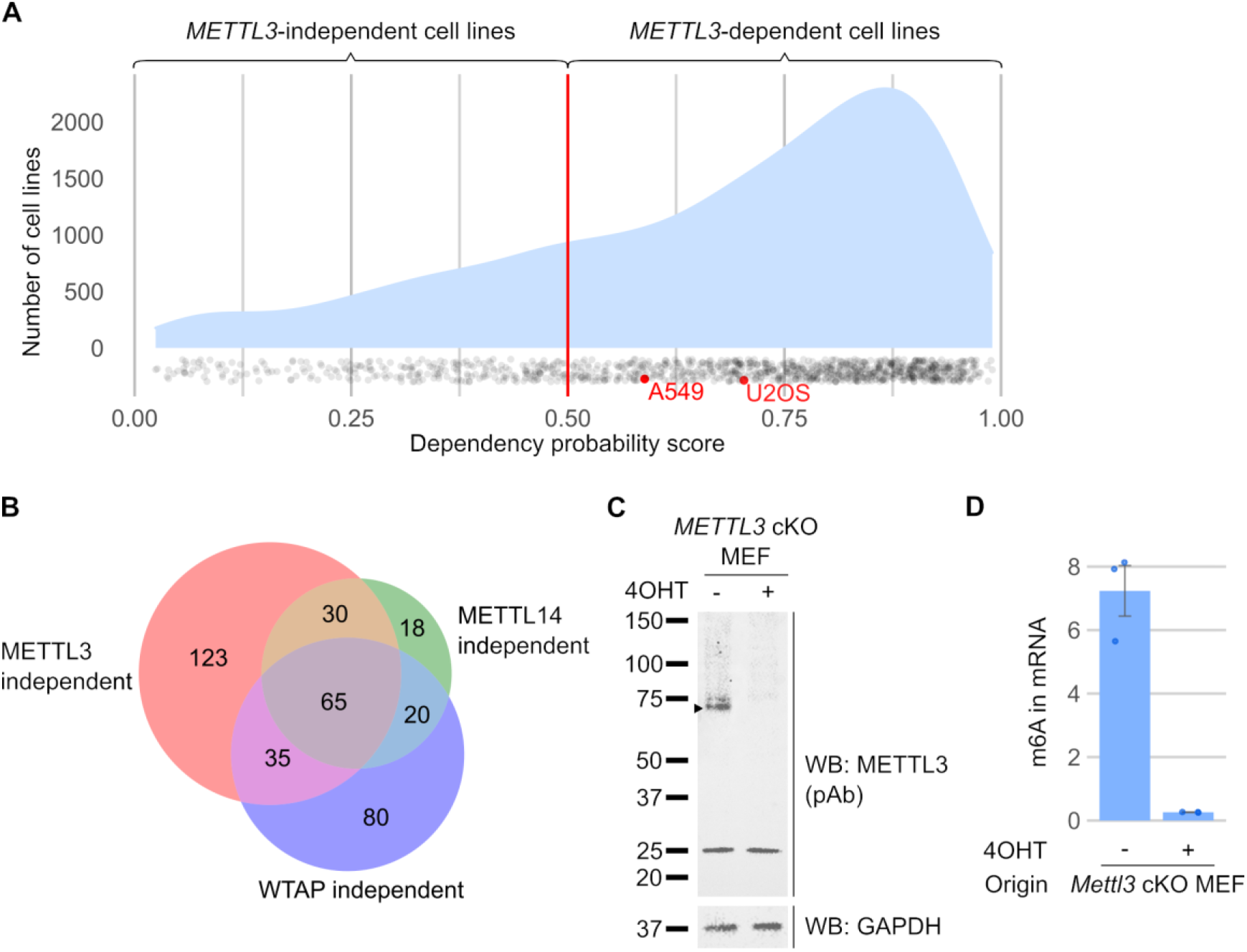
Conditional *METTL3* knockouts can be used to study m^6^A when stable *METTL3* knockouts are not viable. **A** Most cell lines are dependent on *METTL3* for growth. Mouse studies previously indicated that *Mettl3* is an essential gene for early embryonic survival [5], so we wanted to know which cell lines are dependent on *METTL3*. Using the CRISPR gene dependency probability score from the DepMap dataset [40–44], we found that 801 of 1054 cell lines are dependent on *METTL3* (dependency probability score > 0.5). Therefore most cells lines will not survive after *METTL3* knockout. The density plot shows the overall distribution of dependency probability scores, while each individual cell line is represented a dot. U2OS and A549 cell lines (red), where *METTL3* has been previously knocked out, are shown here to be dependent on METTL3 and thus should not be viable after *METTL3* knockout. **B** A small subset of cell lines may be m^6^A-independent. Although most cell lines are dependent on *METTL3*, a small subset of cell lines can survive despite *METTL3* knockout. To identify cell lines which we can confidently consider m^6^A-independent, we obtained a list of cell lines whose survival is also independent of other members of the m^6^A-writer complex, *METTL3, METTL14* and *WTAP*. This approach shows that 65 cell lines can proliferate in an m^6^A-independent manner (**S3 Table**). **C** *Mettl3* conditional knockout MEFs do not express METTL3 protein. To determine the amount of m^6^A in mRNA that can be attributed to *Mettl3*, we generated a tamoxifen-inducible *Mettl3* conditional knockout MEF cell line. We used western blot to validate the loss of METTL3. After 5 days of 4-hydroxytamoxifen treatment (500 nM), we observed loss of the wild-type METTL3 protein. 30 µg per lane. WB = western blot, pAb = polyclonal antibody. **D** *Mettl3* conditional knockout MEFs show near-complete loss of m^6^A. We measured m^6^A levels in mRNA derived from tamoxifen-inducible *Mettl3* conditional knockout MEFs. 8 days after 4-hydroxytamoxifen treatment (500 nM), the *Mettl3* KO MEFs showed 3.6% remaining m^6^A. Hence, m^6^A is almost completely lost after *Mettl3* knockout in MEFs. Error bars indicate standard error (n = 3).

Thus, this leads us to believe that reported stable *METTL3* knockout cell lines were able to survive because they produce a functional alternatively spliced METTL3 isoform that bypasses the CRISPR mutation, similar to the HC *Mettl3* KO mESCs. This alternative METTL3 isoform also explains why these cells retain high levels of m^6^A. Thus, the reported *METTL3* knockout cell lines are likely not true *METTL3* knockouts.

Only a few cell lines are able to survive after *METTL3* knockout. The 21Q4 DepMap dataset shows that only 253 cell lines are not dependent on *METTL3*. Furthermore, only 65 cell lines were not dependent on any of the members of the m^6^A writer complex, which includes *METTL3, METTL14* and *WTAP* [6,21,22,45] (**Fig 4B, S3 Table**). This small subset of cell lines may be the best cell lines for creating *METTL3* knockout cells, since loss of m^6^A in the other cell lines will most likely lead to cell death unless an alternatively spliced METTL3 isoform is selected for. Thus, any reported stable *METTL3* knockout in cell lines other this small subset of m^6^A-independent cell lines should be more carefully evaluated.

### METTL3 is responsible for all m^6^A in mRNA in mouse embryonic fibroblasts

Although the full *Mettl3* knockout in mESCs led to near complete loss of m^6^A [5], this may be a unique feature of embryonic stem cells. It is possible that other cell types can compensate for METTL3 depletion using other methyltransferases that can synthesize m^6^A. To find out if METTL3 is the major writer of m^6^A in mRNA in cell lines other than mESCs, we needed to deplete METTL3 in a different cell line, and measure the remaining amount of m^6^A. Because stable *METTL3* knockout cells are generally not viable, we produced *Mettl3* conditional knockout cell lines derived from *Mettl3*^*fl/fl*^ mice. *loxP* sites flanking exon 4 of METTL3 were inserted in the mouse genome [32]. Upon tamoxifen-induced expression of Cre, recombination of the *loxP* sites leads to genomic deletion of exon 4 which encodes the ZFD, similar to how *Mettl3* is deleted in the JH *Mettl3* KO mESCs.

Thus, we produced an immortalized mouse embryonic fibroblast (MEF) cell line with tamoxifen-inducible *Mettl3* knockout. In tamoxifen-treated MEFs, we found that METTL3 protein expression was completely lost 5 days after tamoxifen-induced *Mettl3* knockout (**Fig 4C, Fig S3A**). By day 6, we observed that MEF proliferation was markedly reduced (**Fig S3B**), corroborating that *METTL3* is essential for proliferation, and therefore stable *METTL3* knockout cell lines are unlikely to survive.

We next determined the fraction of m^6^A in mRNA that is catalyzed by METTL3 in MEFs. We found that 8 days after tamoxifen treatment, most of the m^6^A was lost, with only 3.6% of m^6^A remaining (**Fig 4D**). Thus, this demonstrates that aside from mESCs, METTL3 is also the predominant m^6^A methyltransferase in MEFs.

## DISCUSSION

Here we address the source of METTL3-independent m^6^A, which is widely discussed in the literature and have been described based on high levels of residual m^6^A after knockdown or knockout of *METTL3* [4,7,8,21–32,38,39,45–50]. This has led to the idea that some m^6^A is catalyzed by METTL3, while another substantial fraction of m^6^A is catalyzed by other enzymes. Our results show that the residual m^6^A seen after *METTL3* knockout can be readily explained by the induction of alternatively spliced hypomorphic *METTL3* alleles and the subsequent expression of altered METTL3 proteins. We show that a widely used “*Mettl3* knockout” mESC line [4] undergoes alternative splicing to bypass the CRISPR/Cas9-induced mutations, creating a smaller but catalytically active METTL3 protein. We further show that another published *METTL3* knockout U2OS cell line that also has been shown to contain high levels of residual m^6^A [23,24], also exhibits an altered METTL3 protein. These studies show that altered METTL3 proteins likely account for the residual m^6^A in the cells, rather than an alternative methyltransferase. The evidence for the importance of METTL3 alone as the major m^6^A-forming enzyme is supported by our data in which deleting *METTL3* in MEFs using a large genomic deletion leads to a loss of > 95% of m^6^A. Lastly, we demonstrate that METTL3 is essential for the proliferation and growth of the vast majority of cell lines and therefore, many reported stable *METTL3* knockout cell lines are unlikely to be true knockout cells since they would not be able to proliferate, with a few exceptions. Overall, our studies resolve an important long-standing inconsistency in the literature and argue that (1) METTL3 is the major source of m^6^A in mRNA; and (2) m^6^A which remains after *METTL3* knockout is likely due to cells escaping *METTL3* deletion by creation of new METTL3 isoforms.

Our data suggests that *METTL3* knockout can be validated by measuring residual m^6^A levels in mRNA. We have shown here that several cell lines described to be *METTL3* knockouts still express METTL3 isoforms and have high levels of m^6^A [23–29]. *METTL3* knockout cells which continue to have m^6^A should not be used to determine the function of m^6^A in any biological process. Because these cells retain considerable amounts of m^6^A, they cannot be used to identify pathways and processes that require m^6^A. Furthermore, these cells may express isoforms of METTL3 with unknown functions and properties. These non-physiological METTL3 isoforms may lead to unexpected results, which may be conflated with findings that arise from loss of m^6^A.

To more reliably inactivate *METTL3*, genomic regions encoding critical enzymatic domains should be deleted, rather than simply mutated, since small mutations may be more readily bypassed through alternative splicing events. Furthermore, as *METTL3* knockout is not viable in most cell lines, conditional *METTL3* knockouts provide a useful model for studying m^6^A. METTL3 deletion and m^6^A levels must be carefully documented before reporting these cell lines as *METTL3* knockouts. m^6^A quantification will also allow a better understanding if partial or complete losses of m^6^A are able to mediate the studied outcomes.

Although our results indicate that METTL3 accounts for the vast majority (>95%) of m^6^A in mRNA in MEFs and mESCs, numerous studies have shown that *METTL3* knockout results in a large amount of residual m^6^A in mRNA. Three major technical problems likely account for this “METTL3-independent” m^6^A:

1. Hypomorphic *METTL3* alleles. In this study we characterized a variety of diverse alternative splicing events that occur in *Mettl3* “knockout” mESCs. In these cells, *Mettl3* was targeted using CRISPR/Cas9 systems to create indels and frameshifts. Although this is a reasonable method for gene inactivation [51,52] and may have initially inactivated *Mettl3*, cells which develop alternative splicing patterns that enable formation of an active METTL3 enzyme will gain a proliferative advantage. It is likely that after *METTL3* knockout, cells in which *METTL3* was truly inactivated will stop proliferating. At the same time, any cells that can express a *METTL3* isoform which skips the mutation will gain a proliferative advantage, and will be further selected for their ability to upregulate this isoform. These escaped cells will then be expanded and incorrectly reported as a “*METTL3* knockout” cell line. Similar alternative splicing events that bypass CRISPR/Cas9-induced mutations have been reported in studies of inactivation of other genes [53–55]. Alternative splicing may allow bypass of CRISPR/Cas9-mediated *METTL3* knockout in other cell lines.
2. Heterogeneity of cell population. In a population of *METTL3* knockout cells, any contaminating cells which express functional METTL3 will lead to some level of m^6^A being detected. For example, mouse studies frequently use tissue-specific knockout systems to deplete *Mettl3* in a specific tissue as *Mettl3* knockout is embryonic lethal [5,30–32]. When isolating these tissues, contamination from other cells in the tissue, such as endothelial or immune cells, could lead to detectable m^6^A levels in the samples. Additionally, Cre expression is variable, and can lead to a lack of knockout in some cells [56]. Thus, residual m^6^A in these experiments may simply reflect a mixed population of knockout and non-knockout cells. METTL3 levels can be measured by immunofluorescence to determine if knockout is heterogeneous in these populations.
3. Misattribution of background noise in m^6^A mapping studies as m^6^A sites. A common approach for studying m^6^A sites is MeRIP-seq [3,57], and other antibody-based sequencing methods to detect m^6^A across the transcriptome [58]. In numerous studies, researchers knocked down or deleted METTL3 and regarded the remaining m^6^A “peaks” as “METTL3-independent m^6^A peaks” [50,59]. However, several studies have shown that even in *METTL3* knockout cells where m^6^A cannot be detected, m^6^A peaks can still be observed [60–62]. Thus, these m^6^A peaks should be regarded as background noise, rather than “METTL3-independent m^6^A peaks”. This noise can be due to m^6^A antibodies binding to non-m^6^A sites [58,59,61] or due to nonspecific binding during immunoprecipitation [63]. Thus, peaks that remain after *METTL3* knockout likely do not reflect real m^6^A sites. To identify true METTL3-independent m^6^A, methods such as SCARLET [64] can be used to validate that an m^6^A site in *METTL3* knockout actually reflects m^6^A.

A very small amount of m^6^A clearly remains after *Mettl3* knockout in both the JH *Mettl3* KO mESCs and in our conditional *Mettl3* KO MEFs. An alternative methyltransferase may be able to produce m^6^A on a small number of mRNA transcripts. For example, METTL16, which catalyzes the formation of m^6^A in U6 snRNA, also methylates *MAT2A* mRNA [65,66]. However, the sequence context of METTL16 is CAG [67–69], which differs from the predominant DRACH (D = A/G/U, R = A/G, H = A/C/U) sequence context for METTL3 [58,70,71]. Furthermore, METTL16 methylation is dependent on a hairpin structure found in U6 snRNA and *MAT2A*. Although current attempts to find other METTL16-dependent m^6^A in mRNA have been unsuccessful [59,66], METTL16-dependent m^6^A can likely be recognized based on the CAG sequence context and the U6-like hairpin structural context. m^6^A can also be formed by METTL5-TRMT112 and ZCCHC4 in the 18S and 28S rRNA, respectively [72,73]. Although current attempts have failed to identify m^6^A in mRNA catalyzed by these enzymes [72], a candidate m^6^A mediated by either of these enzymes will likely be in a AAC sequence context as well as the unique structural context utilized by these enzymes [72,74]. Any m^6^A predicted to be “METTL3-independent” is likely to exist within these unique structural contexts and would be lost upon depletion of one of these enzymes.

Importantly, our data does not invalidate the original studies by Batista *et al* on the role of METTL3 in embryonic stem cell differentiation. Batista *et al* used their *Mettl3* hypomorphic mESCs to show that m^6^A is required for differentiation from the naïve pluripotent stem cell state [4]. The study by Geula *et al* using full *Mettl3* knockout mESCs later corroborated this result [5]. The initial study using *Mettl3* hypomorphs is particularly useful as it demonstrates that a partial depletion of m^6^A is sufficient to block differentiation of primed mESCs [5]. Thus, mESCs are highly sensitive to m^6^A levels, and fail to differentiate even with partial loss of m^6^A. This has important implications for researchers attempting to inhibit METTL3 to influence cellular differentiation states, for example in cancer [7,75].

## MATERIALS AND METHODS

### Cell culture

HC *Mettl3* KO and wild-type mESCs were previously described by Batista et al [4], and were a gift from P.J. Batista and H. Chang (Stanford University). JH *Mettl3* KO and wild-type mESCs were previously described by Geula et al [5], and were a gift from S. Geula and J.H. Hanna (Weizmann Institute of Science). All mESCs were grown in gelatinized (0.1% gelatin in water, EmbryoMax ES-006-B) tissue culture plates in mESC media (KnockOut DMEM (Gibco #10829018), 15% heat-inactivated fetal bovine serum (FBS) (Gibco #26140079), 100 U/ml penicillin, 100 µg/ml strepromycin (Gibco #15140122), 1x GlutaMax (Gibco #35050061), 55 µM β-mercaptoethanol (Gibco #21985023), 1x MEM non-essential amino acids (Gibco #11140076), 1000 U/ml LIF (Millipore # ESG1107), 3 µM CHIR99201 (Sigma Aldrich SML1046), 1 µM PD0325901 (APExBIO # A3013)).

*METTL3* KO U2OS cells were previously described by Xiang et al [23], and were a gift from Y. Xiang and Y. Shi (University of Oxford). *METTL3* KO A549 cells were previously described by Courtney et al [38], and were a gift from D.G. Courtney and B.R. Cullen (Duke University Medical Center). U2OS cells and A549 cells were grown in high glucose DMEM (Gibco # 11995073) with 10% FBS (Gibco #26140079) and 1% penicillin-streptomycin (Gibco #15140122).

All cells were grown at 37°C, 5.0% CO2. All cells were passaged as needed using TrypLE Express (Gibco #12604013).

### Western blot

Cells were lysed in RIPA buffer (50 mM Tris-HCl pH 7.5, 200 mM NaCl, 1% NP-40, 0.5% sodium deoxycholate, 0.1% SDS) with 1X Halt protease and phosphatase inhibitor cocktail (Thermo Scientific #78440). Cell debris was cleared by centrifuging at 14000G for 10min. Protein quantification was done by Bradford assay (BioRad #5000201) or Pierce BCA protein assay kit (Thermo Scientific #23225) according to manufacturer’s instructions. Proteins were resuspended in 1X NuPAGE LDS Sample Buffer (Invitrogen NP0007) + 50mM dithiothreitol. Protein samples were separated by electrophoresis using NuPage 4 to 12% Bis-Tris gels (Invitrogen NP0322, NP0335) alongside Precision Plus Protein All Blue Prestained Protein Standards (Bio-Rad #1610373). Proteins were then transferred to nitrocellulose membranes in 1X tris-glycine transfer buffer (25 mM Tris base + 200 mM glycine) + 20% methanol. Membranes were blocked using 5% BSA in phosphate buffered saline (PBS) + 0.1% (v/v) Tween-20 for 1h at room temperature, then stained with primary antibodies overnight. Membranes were then washed with PBS + 0.1% (v/v) Tween-20, then stained with HRP-conjugated secondary antibodies for 1h at room temperature. Membranes were washed with PBS + 0.1% (v/v) Tween-20 to remove excess antibody, and reactive bands were visualized using Pierce ECL Western Blotting Substrate (Thermo Scientific #32109). Western blots images were collected on a BioRad ChemiDoc XRS+ using the Image Lab software (BioRad).

### Antibodies

For immunoblotting experiments, we used anti-METTL3 rabbit polyclonal antibody (ProteinTech #15073-1-AP) directed against amino acids 229 – 580 in METTL3, anti-METTL3 rabbit monoclonal antibody (Cell Signalling Technology #96391S) directed against residues surrounding Leu297 of METTL3, anti-GAPDH rabbit monoclonal antibody (Abcam # ab181603) as primary antibodies. All primary antibodies were diluted 1:1000 in 5% bovine serum albumin in PBS + 0.1% (v/v) Tween-20. Amersham ECL Rabbit IgG, HRP-linked whole antibody from donkey (Cytiva NA934) was used as secondary antibody, at 1:5000 dilution in 5% bovine serum albumin in PBS + 0.1% (v/v) Tween-20.

### 5’ RACE

5’ RACE was performed using the Template Switching RT Enzyme Mix (New England Biolabs M0466) according to manufacturer’s instruction. 1 µg total RNA was incubated with 10mM dNTP and 10mM dT(40) primer at 75°C for 5 min to allow the primer to hybridize. Full-length cDNA was reverse transcribed in 1X template switching RT buffer, 1X template switching RT enzyme, with 3.75 µM template switching oligo (GCT AAT CAT TGC AAG GAT CCG TAT CAA CGC AGA GTA CAT rGrGrG) for 90 min at 42 °C. RNA was degraded using 5 units RNase H (New England Biolabs M0297) for 30 min. 10% of the cDNA product was amplified via PCR using a forward primer against the template switching oligo (CAT TGC AA<u>G GAT CC</u>G TAT CAA C, underlined = BamHI cut site) and a reverse primer against METTL3 exon 6 (CCA GGT AGC G<u>GA TAT C</u>AC AAC, underlined = EcoRV cut site). PCR amplification was performed in 20 uL 1X Phusion® High-Fidelity PCR Master Mix with HF Buffer (NEB M0531) and using touchdown PCR cycling at 98°C for 30s; 5 cycles of 98°C for 10s, 72°C for 60; 5 cycles of 98°C for 10s, 70°C for 60s; 27 cycles of 98°C for 10s, 65°C for 30s, 72°C for 60s; 72°C for 10 min; 4°C hold

The 5’ RACE PCR product was digested using BamHI (New England Biolabs R0136) and EcoRV (New England Biolabs R3195) cut sites found in the primers and cloned into a pcDNA3.1(+) backbone using the Quick Ligation Kit (New England Biolabs M2200). Plasmids were transformed and grown in subcloning efficiency DH5α competent cells (Invitrogen # 18265017) before extraction by Miniprep (QIAGEN #27104) and Sanger sequencing. Sequences of the 5’ RACE products can be found in **S1 Table**.

### Cloning and transfection of METTL3 isoform ORFs

ORFs of METTL3 isoforms were identified from the *Mettl3* mRNAs by finding the longest ORF which encoded an AUG start codon, a protein which aligned with the METTL3 protein, and which was not interrupted by premature stop codons. The full sequence of the METTL3 isoform ORFs identified can be found in **S2 Table**.

The ORFs for full-length METTL3, METTL3-a.i, and METTL3-b.i were constructed via PCR from wild-type *Mettl3* cDNA. FLAG-tags were included at the N-terminus. Primers used for PCR of the METTL3 isoform ORFs can be found in **S4 Table**. PCR amplification was performed in 1X Phusion® High-Fidelity PCR Master Mix with HF Buffer (New England Biolabs M0531) and PCR cycling at 98°C for 30s; 25 cycles of 98°C for 10s, 65°C for 30s, 72°C for 90s; 72°C for 10 min; 4°C hold. The ORFs were then digested using HindIII (New England Biolabs R0104) and BamHI (New England Biolabs R0136), and cloned into a pcDNA3.1(+) vector using the Quick Ligation Kit (New England Biolabs M2200).

Exon-skipping ORFs METTL3-a.ii and METTL3-b.ii were constructed via PCR from wild-type *Mett3* cDNA, followed by assembly and cloning into a pcDNA3.1(+) backbone using Gibson Assembly (New England Biolabs E2611). FLAG-tags were included at the N-terminus. Primers used for PCR of the METTL3 isoform ORFs can be found in **S4 Table**. PCR amplification was performed in 1X Phusion® High-Fidelity PCR Master Mix with HF Buffer (NEB M0531) and PCR cycling at 98°C for 30s; 25 cycles of 98°C for 10s, 65°C for 30s, 72°C for 90s; 72°C for 10 min; 4°C hold. Gibson assembly was performed in 1x Gibson Assembly® Master Mix (New England Biolabs E2611) for 1 hour at 50°C with 1:5 ratio of vector to inserts.

Plasmids were transformed and grown in high efficiency DH5α competent cells (New England Biolabs C2987) before extraction by Miniprep (QIAGEN #27104). Sequences were confirmed via Sanger sequencing.

Plasmids were transfected into mESCs using FuGENE HD transfection reagent (Promega E2311). JH wild-type mESCs or *Mettl3* KO mESCs were plated at 150 000 cells/well on gelatinized (0.1% gelatin in water, EmbryoMax ES-006-B) 6-well plates. The next day, when the mESCs were at 50% confluency, cells were transfected with 2.5 µg of METTL3 isoform ORF-expressing plasmids using 7.5 µl FuGene HD reagent per well. 48 hours after transfection, cell lysate was collected for western blot and RNA was extracted for m^6^A measurements. Successful transfection was confirmed via western blot using anti-FLAG antibody, and anti-METTL3 antibody.

### 2D-TLC measurement of m^6^A

m^6^A levels in mRNA were measured by 2D-TLC (2-Dimensional Thin Layer Chromatography) as described previously [76]. m^6^A was poly-A selected twice using Dynabeads Oligo(dT)25 (Invitrogen 61002) according to manufacturer’s protocol to remove potential contamination from ribosomal RNAs or other non-coding RNAs. 100 ng of twice poly-A selected RNA was then digested by 1 U RNase T1 (Invitrogen AM2283) in 1X PNK buffer for 2h at 37°C. This cuts mRNA after G, therefore only exposing m^6^As in a GA context and omitting m^6^As in the poly-A tail or in other non-mRNA contexts. The 5’ end of the digested RNA is then labelled with 10 U T4 PNK (New England Biolabs M0201) and 10 µCi [γ-^32^P]ATP (Perkin Elmer BLU002Z250UC) for 30 min at 37°C. γ-phosphate was then removed from excess [γ-^32^P]ATP with 0.5 U apyrase (New England Biolabs M0398) in 1X apyrase buffer for 10 min at 30°C. RNA was then digested to single nucleotides using 2 U Nuclease P1 (FUJIFILM Wako Pure Chemical Corporation #145-08221) for 1 hour at 60°C.

2 µl of the digested RNA is spotted on PEI-cellulose TLC plates (Millipore Sigma #105579) 0.5 µl at a time. Plates were developed in 5:3 (v/v) isobutyric acid:0.5 M NH4OH in the first dimension, then in 70:15:15 (v/v/v) isopropanol:water:HCl in the second dimension. Radioactively labelled nucleotides were detected using a phosphor storage screen and Amersham Biosciences Typhoon 9400 Variable Mode Imager. m^6^A levels were quantified using Image Lab software (BioRad).

### DepMap dataset analysis

Gene dependency probability scores were obtained from the DepMap Public 21Q4 CRISPR gene dependency dataset [40–44]. We extracted the gene dependency probability scores of 1054 cell lines on the gene *METTL3* and plotted them to identify cell lines which were dependent on *METTL3*. Gene dependency probability scores greater than 0.5 indicate that a cell line is dependent on the gene.

To identify cell lines which may be independent of m^6^A, we extracted the gene dependency scores of 1054 cell lines on *METTL3, METTL14*, and *WTAP*. We identified cell lines which were independent of each of these genes (gene dependency probability scores < 0.5). We next identified cell lines which were independent of all three genes to obtain a subset of cell lines which may be independent of m^6^A. The full list of predicted m^6^A-independent cell lines can be found in **S3 Table**.

### Generation of *Mettl3* conditional KO MEFs

*Mettl3* conditional knockout mice were previously described by Y. Cheng and M.G. Kharas. *Mettl3*^*flox/flox*^ mice were generated based on construction from the Knockout Mouse Project Repository (KOMP). *loxP* sites were inserted spanning the fourth exon of *Mettl3*.

To generate tamoxifen-inducible *Mettl3* KO MEFs, embryos from *Mettl3*^*flox/flox*^ mice were collected on day 13.5, mechanically separated, trypsinized, and plated. After 3 passages, cells were transduced with SV40 large T-antigen and passaged until they reached senescence. Cells which escaped senescence and became immortalized were then infected with Cre-ERT2 lentivirus. Successfully infected cells were selected with puromycin. Single cell colonies were then isolated and expanded.

To induce the *Mettl3* knockout, MEFs were plated at 100 000 cells/well in a 12-well plate. The next day, when the MEFs were at 50% confluency, cells were treated with 4-hydroxytamoxifen (500 nM) or ethanol (negative control) for 48 hours. Cells continued to be grown and passaged normally after 4-hydroxytamoxifen treatment. Cell lysate was collected five days after tamoxifen treatment for western blot. RNA was collected eight days after tamoxifen treatment for m^6^A measurements.

### MTT assay

To measure the changes in cell proliferation upon *Mettl3* KO, tamoxifen-inducible *Mettl3* KO MEFs were plated at 5 000 cells/well in a 12-well plate. The next day, cells were treated with 4-hydroxytamoxifen (500 nM) or ethanol (negative control). Treatment continued for the duration of the experiment. 0, 2, 4, 6, and 8 days after 4-hydroxytamoxifen treatment, cells were washed with PBS and then incubated in 1:1 (v/v) high-glucose DMEM without phenol red (Gibco 21063029) and MTT (3-(4,5-dimethylthiazol-2-yl)-2,5-diphenyltetrazolium bromide) reagent (5 mg/mL MTT (Abcam ab146345) in PBS) for 3 hours at 37°C. The produced formazan crystals were dissolved in 1.5x volume MTT solvent (4 mM HCl, 0.1% NP-40 in isopropanol) and incubated for 15min at room temperature while shaking on an orbital shaker. Samples without cells were included as background controls. Absorbance was read at 570 nm to estimate the number of cells.

## ACKNOWLEDGEMENTS

We thank members of the Jaffrey Lab for helpful comments and suggestions. This work was supported by NIH grant R35NS111631 to S.R.J, F32CA22104 to B.F.P., Agency for Science, Technology And Research (A*STAR) National Science Scholarship to H.X.P.

## AUTHOR CONTRIBUTIONS

S.R.J. and H.X.P. conceived and designed the experiments. H.X.P. carried out experiments, analyzed data, and prepared figures. B.F.P and A.M. performed m^6^A measurements. S.R.J. and H.X.P. wrote the manuscript with help from all the authors.

