## Supporting Information for "Understanding the source of METTL3-independent m^6^A in mRNA"

**S1 Fig. HC *Mettl3* KO mESCs express altered METTL3 proteins while JH *Mettl3* KO mESCs do not.**

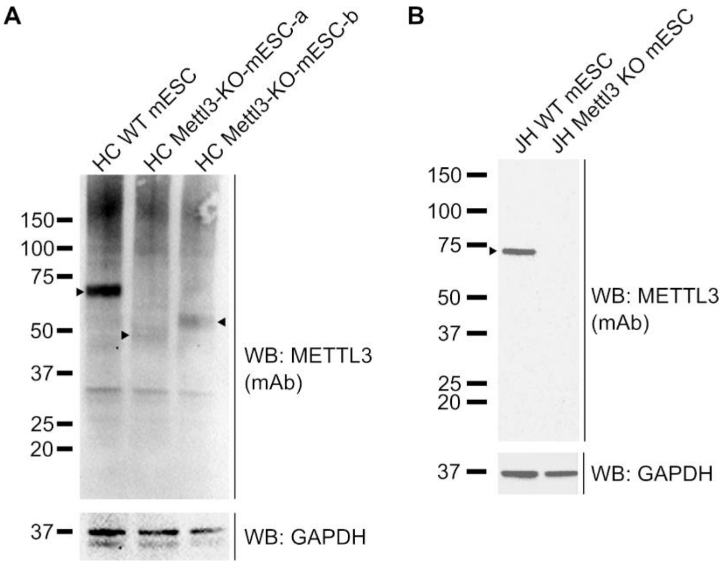


**A** HC *Mettl3* KO mESCs express a novel protein which is immunoreactive to several anti-METTL3 antibodies. We found that HC *Mettl3* KO mESCs express new proteins which are reactive to an anti-METTL3 polyclonal antibody. To confirm that these new proteins may be shortened versions of METTL3, we performed a second western blot with a different anti-METTL3 monoclonal antibody. Similar to the first western blot, we found that full-length METTL3 (75kDa) was lost in both KO cell lines. In the KO cell lines, the same bands which were immunoreactive to the polyclonal anti-METTL3 antibody were also immunoreactive to the anti-METTL3 monoclonal antibody at ~50 kDa in *Mettl3*-KO-mESC-a and ~55kDa in *Mettl3*-KO-mESC-b respectively. This further validates the possibility that the *Mettl3* KO mESCs express smaller versions of METTL3 proteins. 30 µg per lane. WB = western blot, mAb = monoclonal antibody.

**B** JH *Mettl3* KO mESCs do not express METTL3 protein. To investigate the ability of the shortened METTL3 isoforms to synthesize m$^6$A, we needed a cell line which does not express functional METTL3. We used JH *Mettl3* KO mESCs which have no m$^6$A on mRNA. To confirm that JH *Mettl3* KO mESCs do not express METTL3, we performed a second Western blot with a different anti-METTL3 monoclonal antibody. We confirmed that JH *Mettl3* KO mESCs indeed do not express any detectable METTL3 protein. 30 µg per lane. WB = western blot, mAb = monoclonal antibody.

**S2 Fig. *METTL3* KO A549 cells have high m$^6$A levels.**

**A**

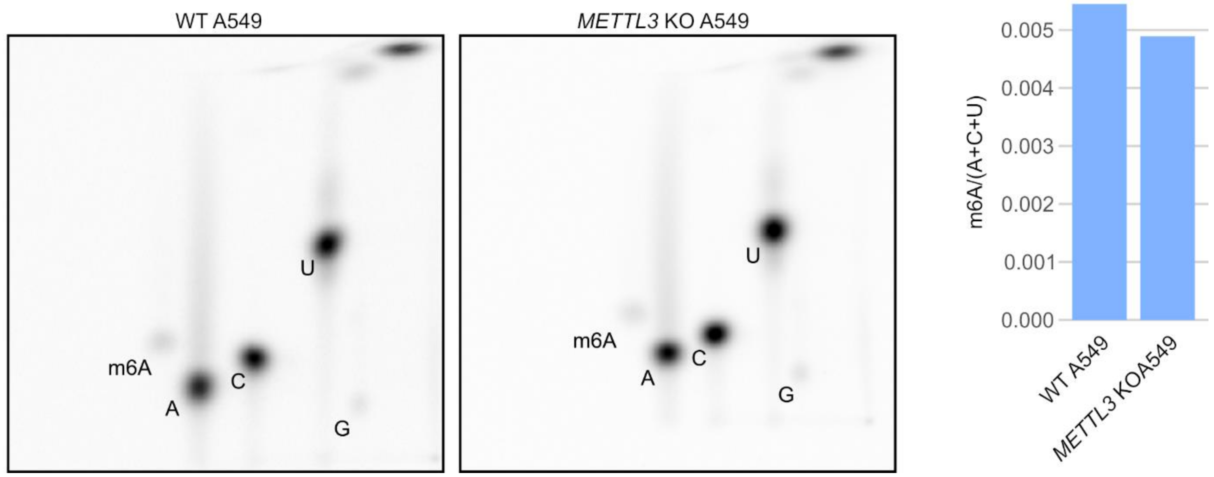


**A** *METTL3* KO A549 cells have persistent m$^6$A. *METTL3* KO A549 cells were previously reported [38]. To find out how much m$^6$A remains in these cells, we measured the levels of m$^6$A using 2D-TLC which measures m$^6$A specifically in the GA context [77]. This limits the detection of m$^6$A to only m$^6$A in mRNAs, where they are found in a DRACH context [58,71,72]. The m$^6$A level in the *METTL3* KO A549 cells was very similar to m$^6$A levels in wild-type A549 cells. This suggests that either the knockout of *METTL3* was incomplete, or that A549 cells may express a non-METTL3 m$^6$A methyltransferase which is responsible for the majority of m$^6$A in mRNAs (n = 1).

**S3 Fig. Depletion of METTL3 leads to slower proliferation in MEFs.**

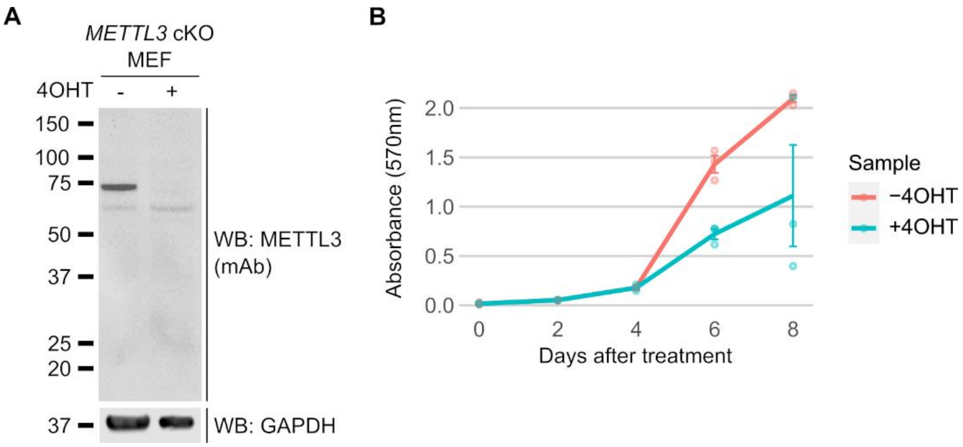


**A** *Mettl3* conditional knockout MEFs do not express METTL3 protein. To confirm that *Mettl3* is the sole m$^6$A writer in cell lines other than mESCs, we produced a tamoxifen-inducible *Mettl3* conditional knockout MEF cell line. We used a Western blot to confirm the loss of METTL3 using a second anti-METTL3 monoclonal antibody. 5 days after 4-hydroxytamoxifen treatment (500nM), we observe loss of the wild-type METTL3 protein. 30 µg per lane. WB = western blot, mAb = monoclonal antibody.

**B** *Mettl3* knockout in MEFs leads to a decrease in cellular proliferation. Using an MTT assay, we measured cell proliferation after 4-hydroxytamoxifen treatment (500nM) over 8 days. Proliferation of *Mettl3* KO MEFs began to slow down compared to wild-type MEFs after 6 days of 4-hydroxytamoxifen treatment. Error bars indicate standard error (n = 3).

**S1 Table. Sequences of 5' RACE products from *Mettl3* in HC *Mettl3* KO mESCs.**

**S2 Table. Sequences of METTL3 ORFs identified in HC *Mettl3* KO mESCs.**

**S3 Table. List of cell lines predicted to be m$^6$A-independent.**

**S4 Table. Primers used in this manuscript.**
